# Comprehensive discovery of subsample gene expression components by information explanation: therapeutic implications in cancer

**DOI:** 10.1101/043257

**Authors:** S. Pepke, G. Ver Steeg

**Affiliations:** Lyrid LLC, South Pasadena, CA; Information Sciences Institute, University of Southern California

**Keywords:** ovarian cancer, RNA-seq, transcriptome, expression components, information optimization, latent factors, machine learning, personalized medicine

## Abstract

**Background:** *De novo* inference of clinically relevant gene function relationships from tumor RNA-seq remains a challenging task. Current methods typically either partition patient samples into a few subtypes or rely upon analysis of pairwise gene correlations (co-expression) that will miss some groups in noisy data. Leveraging higher dimensional information can be expected to increase the power to discern targetable pathways, but this is commonly thought to be an intractable computational problem.

**Methods:** In this work we adapt a recently developed machine learning algorithm, CorEx, that efficiently optimizes over multivariate mutual information for sensitive detection of complex gene relationships. The algorithm can be iteratively applied to generate a hierarchy of latent factors. Patients are stratified relative to each factor and combinatoric survival analyses are performed and interpreted in the context of biological function annotations and protein network interactions that might be utilized to match patients to multiple therapies.

**Results:** Analysis of ovarian tumor RNA-seq samples demonstrates the algorithm’s power to infer well over one hundred biologically interpretable gene cohorts, several times more than standard methods such as hierarchical clustering and k-means. The CorEx factor hierarchy is also informative, with related but distinct gene clusters grouped by upper nodes. Some latent factors correlate with patient survival, including one for a pathway connected with the epithelial-mesenchymal transition in breast cancer that is regulated by a potentially druggable microRNA. Further, combinations of factors lead to a synergistic survival advantage in some cases.

**Conclusions:** In contrast to studies that attempt to partition patients into a small number of subtypes (typically 4 or fewer) for treatment purposes, our approach utilizes subgroup information for combinatoric transcriptional phenotyping. Considering only the 66 gene expression groups that are both found to have significant Gene Ontology enrichment and are small enough to indicate specific drug targets implies a computational phenotype for ovarian cancer that allows for 3^66^ possible patient profiles, enabling truly personalized treatment. The findings here demonstrate a new technique that sheds light on the complexity of gene expression dependencies in tumors and could eventually enable the use of patient RNA-seq profiles for selection of personalized and effective cancer treatments.

## Background

In recent years, innovations in high-throughput sequence-based assays that allow for the rapid and cheap interrogation of the genomes and transcriptomes of individual tumors have given rise to fresh hope for significant progress toward understanding of complex cancer biology and effective treatments [1]. Inference of clinically relevant gene expression networks based upon RNA-seq profiles could provide crucial information in this context. The tumor transcriptome has the potential to provide a view of the functional network alterations that result from a variety of causes, including genetic mutations, copy number variations, epigenetic states, the tumor microenvironment, immunophenotype variations and even transient dynamic effects. In principle, machine learning could be used to leverage the rich information that gene expression provides about each individual to infer tumor-specific features that may guide treatment decisions. Unfortunately, several challenges limit the usefulness of existing methods. The data is high dimensional and the number of samples is small. For this reason, many methods focus on simple linear, pairwise relationships, limiting their generality. Clustering techniques typically use too broad a brush, pigeon-holing patients into a small number of clusters when understanding the interplay of multiple independent factors is crucial. Finally, interpretability of results is essential in this context in order to provide insight into biological mechanisms that could motivate new treatments.

To overcome these limitations, we apply a recently developed machine learning algorithm, CorEx [2, 3], to the analysis of RNA-seq transcriptomes. Multivariate and nonlinear dependencies in the data are captured by total correlation, a generalization of mutual information to multiple variables. Intuitively, CorEx uses an optimization scheme in order to efficiently infer latent variables that explain as much of the dependence in the data as possible. Each latent factor discovered by CorEx is a function of some (possibly overlapping) subset of the input variables (genes) with high total multivariate information, and reflects a relatively independent axis of variation in comparison to the other latent factors. Further, each latent factor defines a separate probabilistic clustering of samples, and thus provides a factor-specific stratification of tumors according to expression patterns of subgroups of genes across the tumor samples. We can apply CorEx again to these inferred latent factors to discover a hierarchy of dependencies among genes.

Ovarian cancer is the deadliest gynecologic malignancy due to its typically late stage at diagnosis, however it is responsive to a wide variety of therapies. In addition to the standard platinum/taxane combination for frontline, active second line therapies include pegylated doxorubicin, topoisomerase inhibitors, angiogenesis inhibitors, anti-metabolites and PARP inhibitors [4, 5]. Ovarian cancer also appears to be an immunogenic tumor for which checkpoint inhibitors, vaccines, and other immunotherapies may provide meaningful benefit [6, 7, 8, 9]. Thus ovarian cancer tumors can be expected to display a rich variety of gene expression patterns related to outcomes under various treatments. On the other hand, current second line treatments are effective for less than 30% of patients, are not curative, and come with often serious cumulative toxicities. With the exception of DNA repair pathway mutations for PARP inhibitors, there are no genetic markers currently in use to predict therapeutic response. Thus new ways to match ovarian cancer patients to effective therapies based upon individual tumor biology will likely yield substantial gains in survival.

In this work we show how the use of CorEx to analyze ovarian cancer RNA-seq profiles uncovers relatively small cohorts of genes that exhibit coherent patterns of differential expression among tumors and that, in many cases, these cohorts are associated with biological pathways as indicated by database annotations of the genes. The lower levels of the hierarchy of inferred gene clusters provide a parsing of the upper layer nodes with respect to function. An example is a layer one node that groups together regulatory noncoding RNAs to regulated protein coding elements. Patient stratification with respect to individual gene groups shows differential survival associations in groups associated with known factors in chemo-responsiveness. Even more significantly, stratifications based upon combinations of group factors display synergistic survival associations. We highlight discovery of a component containing genes related to epigenetic control of stem cell characteristics that have not been previously observed in ovarian tumors. Additionally, when breast cancer data not used for training is stratified relative to this latent factor, a survival association is also observed. Finally, we compare the group results in ovarian tumors to expression profiles in normal ovarian tissue and demonstrate substantial differences in some cases. The strong results presented here demonstrate a powerful tool for dissecting therapeutically relevant gene expression relationships in tumor RNA-seq, and suggest it may be usefully applied to infer complex correlation structure from other types of high-dimensional, noisy biological data [10].

## Methods

### CorEx Algorithm

The random variables, *X*_1_,…, *X*_*n*_, written as *X* for short, represent measured gene expression values. Our goal is to learn a small number of latent factors, *Y* = *Y*_1_,…, *Y*_*m*_, so that the *X*_*i*_’s are (nearly) independent after conditioning on *Y*. In other words, we are looking for latent factors that explain the relationships in the gene expression data. Each latent factor will depend on some subset of the genes, and it can take discrete values, *Y*_*j*_ = 1,…,*c*, that correspond to relative expression levels for that group of genes. We found that stratifying each latent factor into three groups was adequate for capturing relationships without causing problems with estimation. Therefore, our procedure defines a clustering of genes associated with each *Y*_*j*_ and, for each group of genes, a stratification of patients into three groups. Furthermore, this clustering procedure can be iteratively applied to the latent factors, *Y*, themselves to produce a hierarchical clustering of the genes. We now review the technical approach for discovering the latent factors and clusters.

We assume that the measured expression profile for one patient corresponds to a sample, *x*, drawn from some unknown distribution over the random variables (genes), *X*, which is written as *p*(*X* = *x*). Similarly, we can write the marginal probability for a single gene as *p*(*X*_*i*_ = *x*_*i*_). Multivariate mutual information, or total correlation (TC), quantifies the amount of dependence among a set of variables [11]. Total correlation can be defined in terms of the Kullback-Leibler divergence, *D*_*KL*_, or in terms of *H*, the Shannon entropy [12] as follows.

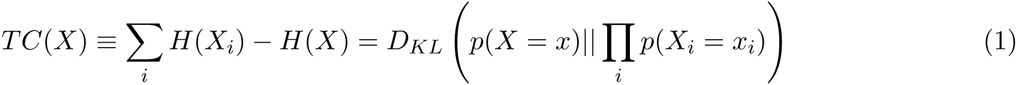

Intuitively, TC represents the distance between the true distribution, and the distribution under the null hypothesis that all variables are independent. TC is zero if and only if the variables are all independent^1^. If we could enumerate all of the underlying causes of dependence in the data, then the TC conditioned on these factors would be exactly zero.

The recently introduced method of Total Correlation Explanation (CorEx) is an information-theoretic optimization that reconstructs latent factors that explain as much of the dependence in the data as possible [2, 3]. The principle behind CorEx is to search for a small set of latent factors, *Y* = *Y*_1_,…, *Y*_*m*_, so that the TC among the original variables is minimized after conditioning on *Y*. We refer to *TC*(*X*; *Y*) = *TC*(*X*)*−TC*(*X|Y*) as the correlation in *X* that is explained by *Y* and we seek to maximize this expression.

We would like to maximize *TC*(*X*; *Y*) over all ways of assigning discrete values to each *Y*_*j*_ as a function of *X*. As mentioned, we restrict *Y*_*j*_’s to take three possible values so that, e.g., *Y*_*j*_ = 0, 1, and 2 might correspond to relatively low, neutral, and high expression. For each sample of data, *X* = *x*^*l*^,with *l* = 1,…, *N* indexing the samples, we define a probability distribution over possible values for each latent factor *j*, *p*(*Y*_*j*_ = *y*_*j*_*|X* = *x*^*l*^). Surprisingly, the optimization of *TC*(*X*; *Y*) over all of these probability distributions is computationally efficient and has low sample complexity [3]. We now describe the iterative procedure for solving this optimization.

What makes our optimization tractable is that we can write the solution for *p*(*Y*_*j*_ = *y*_*j*_*|X* = *x*) in an analytic form that defines an iterative update scheme that is guaranteed to converge to a local optimum. We begin at iteration *t* = 0, with a random probability distribution, *p*^*t*=0^(*Y*_*j*_ = *y*_*j*_*|X* = *x*^*l*^), over each latent factor and for each sample. The solution depends on two types of parameters. First, *α*_*i,j*_ are weights between zero and one that determine how much *Y*_*j*_ depends on *X*_*i*_. Second, *p*(*X*_*i*_ = *x*_*i*_*|Y*_*j*_ = *c*) is the marginal probability of observing *X*_*i*_ = *x*_*i*_ given an observed label *Y*_*j*_ = *c*. We take this distribution to be a normal distribution with mean, *µ_i,j,c_* and variance, *σ_i,j,c_*. The parameters, *α*_*i,j*_, *µ*_*i,j,c,*_ *σ*_*i,j,c*_, along with the class weights, can be updated at each time *t* according to the following rules:

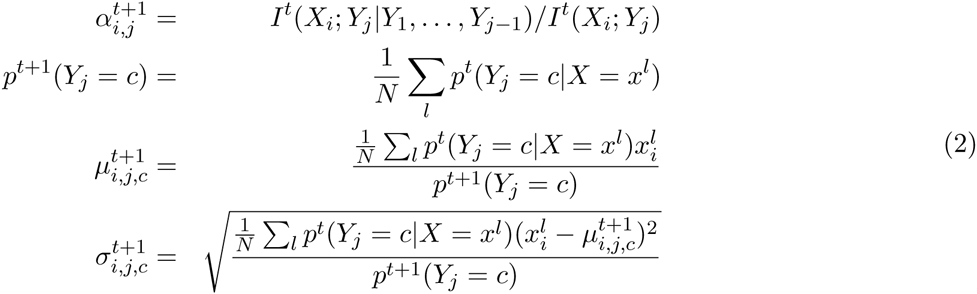

In practice, we alter the first rule to speed up computation and we devised a novel Bayesian estimate for the last two update rules that improves results on datasets with a small number of samples. Details of these revised updates are described in Supplementary Methods.

Once we have estimates at time step *t* of 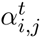 and *p*^*t*^(*X*_*i*_ = *x*_*i*_*|Y*_*j*_ = *y*_*j*_), then we can update the distribution of latent factors to get a new solution that is guaranteed to increase our objective. The solution has the following form.

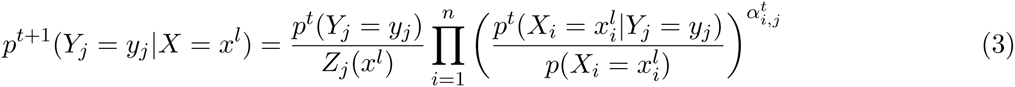

In this expression, *Z*_*j*_(*x*) is a normalization constant. It is fortunate that this expression depends only on marginal distributions between *X*_*i*_ and *Y*_*j*_ because these can be estimated accurately even with small amounts of data. Furthermore, there is no dependence on the pairwise correlations among the *X*_*i*_, so that the overall computational complexity is linear in the number of variables while most comparable methods (like PCA or hierarchical clustering) are at least quadratic. Now that we have update rules for the probabilistic labels, and for the parameters, we simply iterate between them until we achieve convergence.

The scheme described finds a set of latent factors, *Y*, that explain as much of the dependence in *X* as possible. These latent factors, *Y*, are also guaranteed to be less correlated than the original variables, *X*, as measured by TC [3]. Therefore, we can also view the latent factors as approximately recovering independent components. In results below, we consider how combinations of these quasi-independent factors can improve our ability to predict long-term health outcomes.

If the latent factors are not completely independent, we can repeat the procedure we have just described to learn a second layer of latent factors, say *Y* ^2^, that explain the dependence in the first layer, *Y* ^1^’s, by maximizing *TC*(*Y* ^1^; *Y* ^2^). We will continue until the latent factors at layer *k* are completely independent so that *TC*(*Y* ^*k*^) = 0, which implies that, for any *Y* ^*k*+1^, *TC*(*Y* ^*k*^; *Y* ^*k*+1^) ≤ *TC*(*Y* ^*k*^) = 0. In other words, our optimization will find that there is no dependence left to explain. The resulting hierarchy of relationships is compared to standard hierarchical clustering in the Results section. Note that at at the higher levels of the hierarchy, both the inputs and latent factors are discrete, leading to a simpler update scheme described in the Supplementary Material.

The quality of the solution that is found by CorEx is quantified through the expression *TC*(*X*; *Y*). As we add more latent factors this objective increases before achieving a peak and then decreasing. Because of computational limitations, we used only 200 latent factors at the first layer even though more would have resulted in higher TC. At the higher levels, we tried different numbers of latent factors to optimize *TC*(*X*; *Y*) leading us to use 30 factors at layer 2 and 8 at layer 3. With this number of factors, each CorEx run took about two days on a single Amazon “r3.xlarge” node with 30 GB of RAM.

As a clustering method, CorEx has been favorably compared against standard methods like spectral clustering, k-means, and hierarchical clustering [2]. CorEx can also be viewed as a type of dimensionality reduction and in that capacity has been compared to PCA, independent component analysis, non-negative matrix factorization, and Isomap [2]. More generally, CorEx represents a new approach to unsupervised deep representation learning and, as such, is best compared to “neural network” approaches like autoen-coders and restricted Boltzmann machines against which it has demonstrated the ability to recover much more interpretable structure [3].

### Data Sources

Gene level RNA-seq values (normalized to reads per kilobase per million or RPKM) for 420 high grade serous ovarian tumor samples and 780 breast cancer samples were downloaded from the TCGA data portal (The TCGA Research Network: http://cancergenome.nih.gov), along with available clinical metadata for the samples. Somatic mutation data for 316 samples was obtained separately from the cBioPortal for Cancer Genomics [13, 14]. A training set of genes was selected by combining the RPKM data for the top 3000 genes by variance in expression with the RPKM values for the 2371 genes that were mutated in more than one sample. The gene RPKM values for the selected genes were then converted to z-scores to form the 420x5371 training matrix. 376 samples were associated with survival times in the clinical Biotab file and used for the Cox proportional hazard analyses. GTEx normal tissue data [15] for 39 samples was downloaded from the GTEx portal (www.gtexportal.org) and normalized similarly to the tumor data.

### Database Annotations

KEGG, GO, and PPI enrichments for the gene groups were obtained using the stringdb package in R [16, 17]. A maximum of 400 genes were used from each group. For groups from CorEx runs that utilized the Bayes shrinkage prior, only genes with a weighted mutual information greater than a threshold of 0.002 were retained for annotation enrichment purposes. FDR values with respect to the 200 groups (corresponding to the p-value cutoffs used in the Circos [18] plot in Figure 2) were calculated by random selection from the set of 5371 genes used for training. The genes were alloted to groups with a size distribution identical to that of the CorEx discovered groups. In order to obtain summary GO terms for upper layer nodes in the CorEx factor hierarchy, the lists of GO terms from stringdb were submitted to the Revigo web server [19], which clusters GO terms to select a subset of distinct representative terms. GO term information content was approximated as *−log*(*n*_*d*_*/n*_*r*_) where *n*_*d*_ denotes the number of descendants of the term in the gene ontology and *n*_*r*_ is the number of descents of the root term. Hierachical and k-means clustering as well as PCA analysis were done using Cluster 3.0 [20].

**Figure 1.**
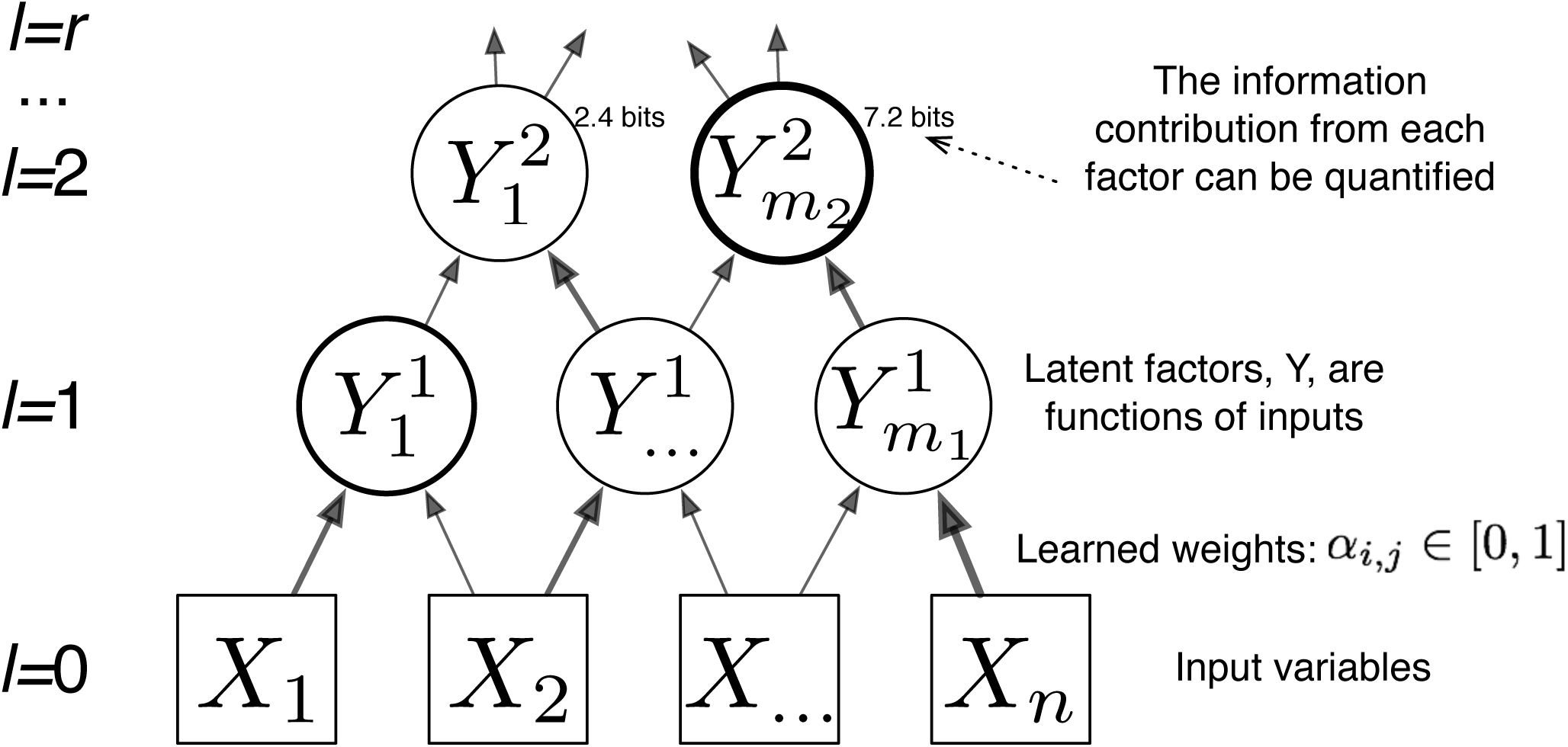
A simple hierarchical model constructed by CorEx. Variables in each layer are associated with latent factors in the next higher layer with a weight that is determined by optimizing over its relative contribution to the explanatory latent factors. Visually we set the thickness of links based on mutual information between variables and the thickness of nodes is proportional to the information contributed by that factor toward the objective, TC(X;Y).

**Figure 2.**
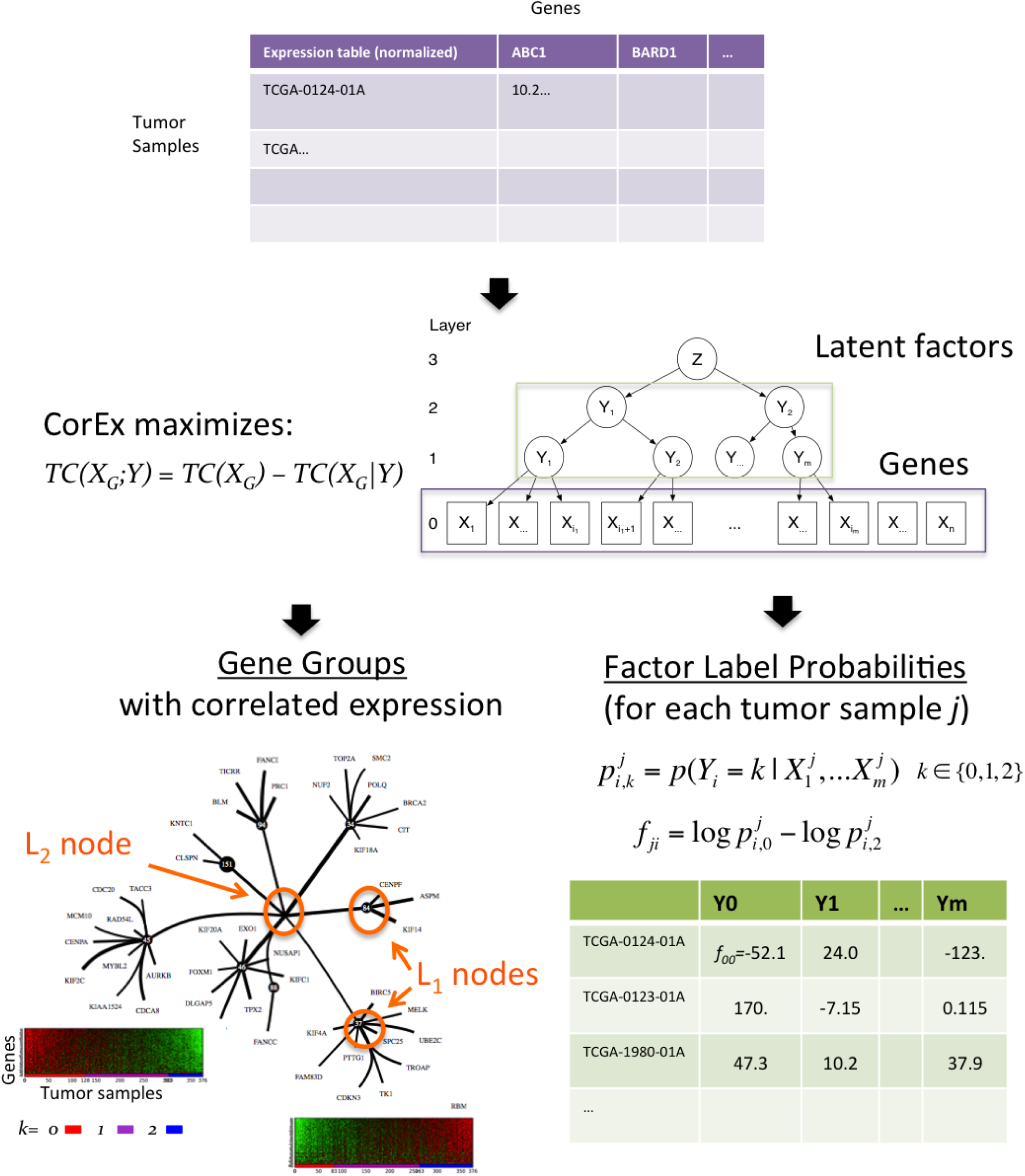
Correlation Explanation algorithm applied to tumor RNA-seq. For training, CorEx is provided only a matrix of normalized gene expression values for the available tumor samples. The number of possible labels for each latent factor is specified, here set to three. In this application, we also set the number of layer one latent factors to 200. CorEx finds probabilistic assignments of genes to latent factors by maximizing the total correlation of the genes in groups, simultaneously minimizing dependence between latent factor groups. The factor labels from lower layers are used as input to upper layers in order to generate a hierarchical model. The output from CorEx is thus a hierarchical model, specified as a set of probabilities characterizing the association of genes or factors with latent factors at the next highest layer as well as probabilities that a given tumor sample?s expression pattern can be explained by each factor in a particular label state. The three probabilities for a given tumor sample and factor are can be usefully summarized by a single value that is the natural logarithm of the probability difference for the factor labels corresponding to extremal expression. The genes with high mutual information relative to latent factors show clear patterns of correlation when viewed on expression heat maps with tumor samples ordered by the summary latent factor score.

### Survival Analysis

For survival analysis, the gene groups were fixsltered to retain only those with fewer than 100 correlated genes in each group above threshold significance and those associated with gene ontology terms with adjusted p-values less than a 1% FDR threshold. Additionally, fourteen groups were eliminated that were presumed to represent chromosomal location correlations rather than network interactions. This filtering resulted in 66 groups that were then analyzed for single factor survival associations under a coxph model in R. Patients were stratified according to relative risk under a predictive coxph model. Patients were assigned to one of three groups according to whether they were in the bottom 30%, top 30%, or middle 40% of risk of death. The top individual factors with differences in survival (determined by p-values between the two extremal strata as calculated by the survdiff function less than 0.05) were retained for combination analysis. The false discovery rate for this procedure was estimated by repeatedly selecting 66 factor groups (200 times), randomly shuffling their factor labels with respect to patient time of death and calculating p-values for both single and combination factors in the same fashion as before. False discovery rates were then calculated as the number of randomized groups or combinations exhibiting p-values less than a given threshold divided by the observed number of groups or combinations with p-values less than that threshold.

## Results

### CorEx infers gene expression relationships from tumor RNA-seq profiles

CorEx can be used to construct a hierarchy of latent factors that “explain” correlated gene expression among samples with respect to subsets of genes (Figure 2). Details of the algorithm and implementation can be found in the Methods section and the references therein. In the context of RNA-seq tumor profiles, the input to the algorithm is a training matrix consisting of z-scores for each gene among all the samples. The CorEx output is a set of *m* gene groups with the genes ordered within a group according to their estimated mutual information (MI) with respect to the group’s latent factor and a probabilistic assignment of samples to the *k* factor labels. Thus for each sample *j*, group *i*, and latent factor label *k* (here *k* ∈ {0, 1, 2}), CorEx provides an estimate of 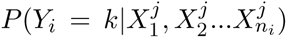, where 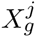 denotes the expression of gene *g* in the *j*^*th*^ sample. Here the *P* (*Y* = *k|X*_1_, *X*_2_*…X*_*n*_) for each sample is converted to a single score, *f*_*ji*_ that indicates sample *j*’s congruence with the factor *i* label values representing either very high or very low gene expression. This score provides an ordering of the samples with respect to gene expression. Heat maps of genes within groups ordered by MI versus samples ordered by *f*_*ji*_ show how groups with the greatest TC display strong correlations among many genes, with the clearest relationships being between genes with high group MI, whereas lower TC groups show noisier correlations and/or few genes (Figure 2 and Supplementary Fig 1 of heat maps, and the Supplementary data files).

Sharing of genes among groups is an integral part of the algorithm and this gene sharing is desirable for a couple of reasons. For one, it known that gene products often perform multiple roles within the cell and therefore may participate in multiple interaction networks. For another, given that TC increases with the number of genes (all other things being equal), it is easier for the algorithm to find very small expression cohorts that participate in larger networks when they can be manifested as similar large network groups with different sub-clusters ranked most highly. In this application, genes tend to be associated with several groups on average, however typically only one to four at relatively high MI values.

Due to the complexity of the search space, the same gene groupings are not guaranteed to be found in different runs. However, comparison of gene lists between runs demonstrates that groups with high TC per gene are very likely to be reproduced. Additionally, we find that a Bayes shrinkage prior over the factor probabilities increases reproducibility of groups with low TC, at the cost of possibly suppressing some real but weak signals (Supplementary Figure 3). In this work we fix the number of latent factors in layer one to 200. Though this may seem like a relatively large number, the vast majority CorEx factors found appear to contain meaningful correlations. This is evident from a comparison of the group TC values to those resulting from training on a randomly permuted expression matrix (Supplementary Figure 4). The large TC values relative to the randomized data strongly suggest that the discovered gene groups are not merely the result of chance correlations in the data. As one moves down the list of groups according to total correlation, confidence in the clusters decreases though there appear to be still biologically meaningful clusters at the lower end, suggesting that training on a greater number of samples or taking into account additional biological information will be useful. The latter is investigated in the following sections.

### CorEx gene groups are enriched for protein-protein interactions, GO, and KEGG annotations

A majority of the CorEx gene groups show enrichment for genes that function within specific biological pathways and/or networks as indicated by significant protein-protein interactions (PPI), Gene Ontology terms [21], and KEGG [22] pathway annotations (summarized in Figure 3a). Multiple groups are found related to the mitotic cell cycle and its regulation, extracellular matrix organization and interaction, chromatin modification and DNA packaging, electron transport, and regulation of map kinase and growth factor pathways, among others. Genes that code for proteins that participate in larger functional complexes such as ribosomes are also grouped together. Some of the particular PPI graphs are shown in Figure 3 where it can be seen that CorEx finds networks of genes related to particular immune processes such as Type I interferon signaling, antigen processing and presentation, immune cell activation, migration, and aggregation. In fact, a plurality of found groups have predominantly immune system-related annotations. Overall, the number of groups enriched for annotations as well as their diversity and specificity support the usefulness of CorEx for parsing the information about differentially activated biological processes from the RNA-seq profiles at an unprecedented level of detail. (See Supplementary Figure 2 and Supplementary data for all interaction graphs and listing of terms.)

**Figure 3.**
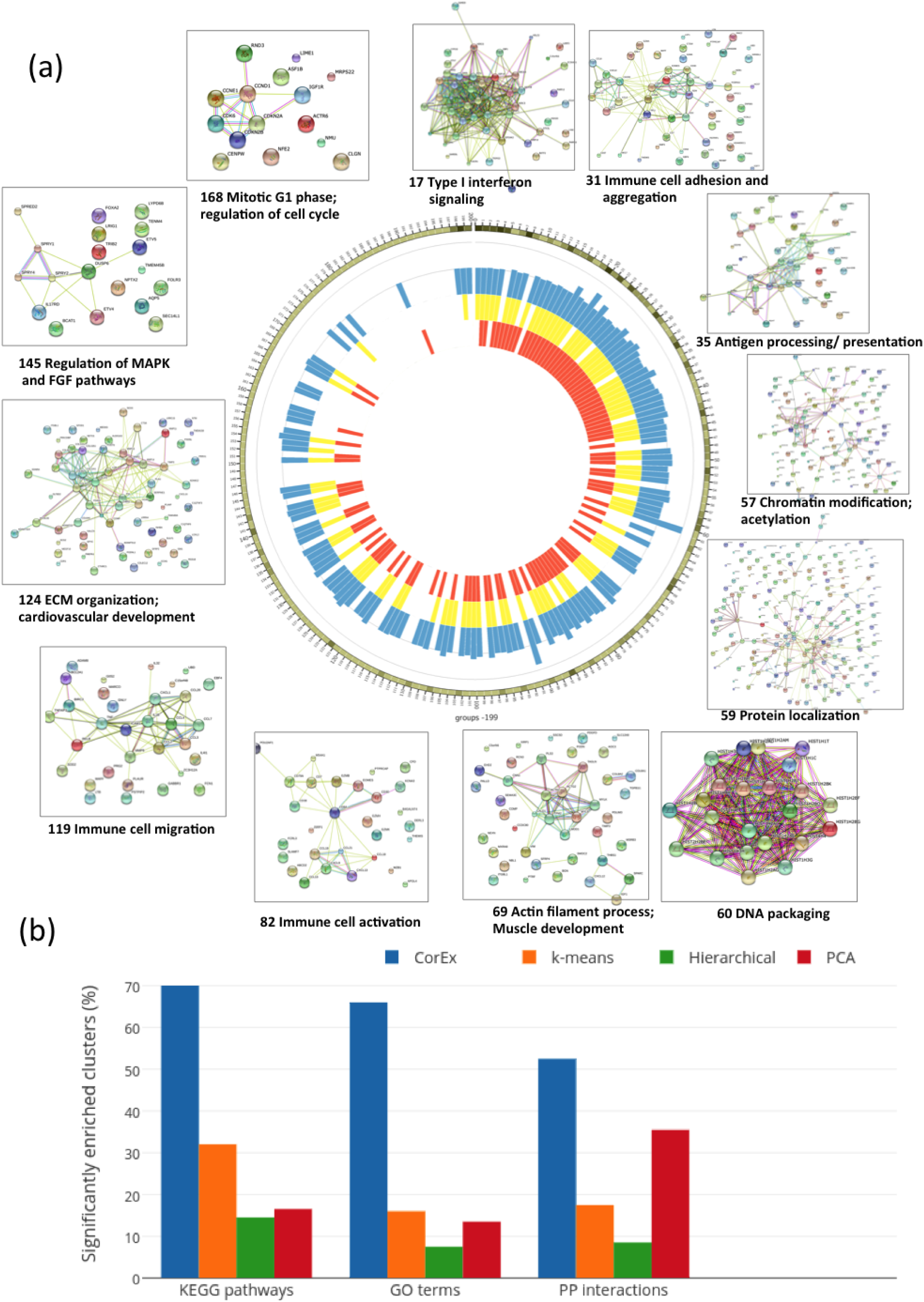
Gene Ontology, KEGG, and protein interaction enrichment for CorEx gene groups. A circos plot indicates which latent factor groups of the 200 inferred by CorEx are enriched for genes on KEGG pathways (yellow), Gene Ontology terms (orange) and Protein-Protein interactions (blue). A tick mark of the corresponding color is drawn for groups exhibiting enrichment greater than a threshold. Further, the blue PPI tick marks have heights proportional to the relative enrichment (observed/expected). The group results appear in order of decreasing Total Correlation, with the number of genes in a group indicated by the green shading on the outer ring of the circos plot. (darker means a greater number of genes with a range from 4 to 1135) Cutoffs were initially chosen according to the presence of coherent biological signals, however randomization tests indicate corresponding FDR values of 1%, 5%, and 10% for GO, KEGG, and PPI, respectively. The vast majority of CorEx groups show annotation enrichment, typically for all three types. A sample of string network connections are shown on the outside, showing both a diversity of network function and relatively few apparently spuriously associated genes within the groups.

The CorEx groups often display especially clear PPI enrichments with relatively little noise, though all three types of annotations typically exceed threshold together. Groups with greater total correlation are more likely to yield significant database annotations. However, because groups with high TC tend to be larger, groups with somewhat lower TC tend have more specific functional annotations at low p-values. It can be seen that many groups even with quite low TC sometimes have associated annotations. Capturing these groups with relatively weak signals that nonetheless appear biologically coherent was part of the motivation for specifying such a large number of factors. The database annotations offer some guidance as to which may merit closer examination. While some enrichment of annotations within the gene groups is not surprising, CorEx substantially outperforms other methods such as hierarchical clustering, k-means, and principal component analysis in terms of the percentage of gene groups associated with significant biological annotations (Figure 3b).

Some of the power of CorEx in this context comes from lifting the restriction of considering only pairwise gene interactions, common in other approaches [23, 24, 25]. As a concrete example of the impact of this less restrictive inference method, one need only examine the pairwise relationships of the top ranked genes in one of the CorEx groups, pictured in Figure 4. The colors of the scatter points indicate the CorEx label value for each tumor sample. In the figure it is clearly evident that the top genes have only weak pairwise expression correlations, yet the clouds of similarly hued points remain coherent across the various plots showing their higher dimensional correlation. The corresponding expression heat map for group 106 (Supplementary Figure 1) also shows clearly that, while any individual gene appears somewhat noisy, the trend across the whole set is easily discernible. The full set of genes from this group exhibit PPI enrichment as well as significant enrichment for the KEGG pathways: ‘pathways in cancer’, ‘Wnt signaling pathway’, ‘proteoglycans in cancer’, ‘basal cell carcinoma’, and ‘Hippo signaling pathway’. CorEx detects this important set of expression relationships that would almost certainly be overlooked based upon pairwise correlations. Additionally, the sensitivity for groups that exhibit stronger pairwise correlation for the top genes can be increased when those signals are reinforced with signals from lower-ranked genes exhibiting weak pairwise correlations but high TC relative to the group as a whole.

**Figure 4.**
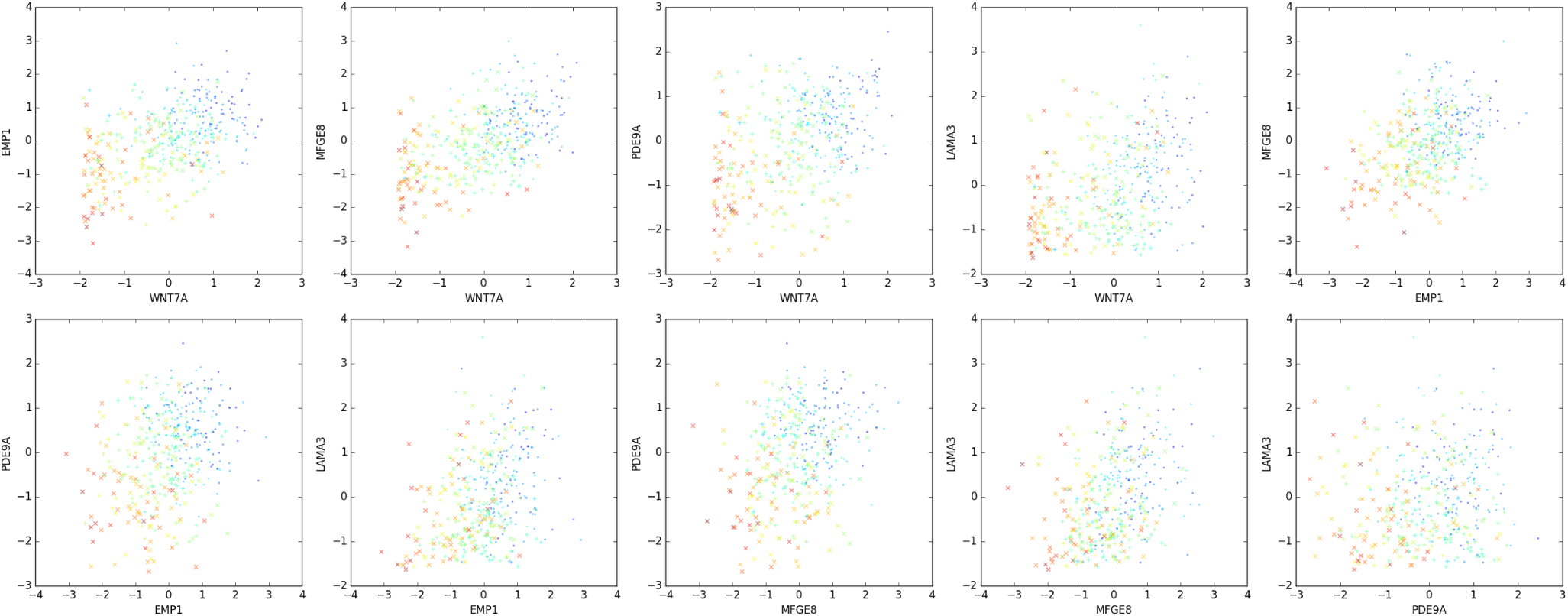
The use of multivariate mutual information increases the clustering power of CorEx. Pairwise correlations of normalized expression of individual samples for the top genes in group 106 appear very weak. The factor label values (indicated by color) show strong coherence across the different plots indicating how samples cluster more strongly in terms of total correlation.

### The CorEx hierarchy of factors has biological significance

The hierarchy of CorEx latent factors appears to reflect biological organization on multiple levels. Though the leaves in the CorEx tree have a great diversity of associated GO terms, those that are shared between children and their common parent node display a trend toward increased enrichment of higher level terms in the L1 cluster (Supplementary Figure 5). Some of these higher level functional relationships are highlighted in Figure 5. Clustering of Gene Ontology terms in the groups reveal layer one and higher clusters for immune signaling, extra-cellular matrix organization, immune cell activation, mitotic cell cycle, microtubule-related processes, and electron transport/chromosome organization. While in some cases this is due to redundancy of membership among the layer zero gene groups (though the genes are ranked differently within the groups, the GO analysis here did not account for relative importance), in others there is a clear distinction in gene membership and function on the finer scale. For example, both groups 31 and 110 are incorporated into the large immune-related cluster near the center of Figure 5a. However, these groups have both different associated GO annotations and zero genes in common. Group 31 has gene ontology terms associated with immune cell activation (e.g. GO:45321 for genes EBI3, Cd79A, BATF, FCER1G, CCL5, EOMES, CD3D, Cd7, LCP1, ZNF683, CD2, ITK, THEMIS, B2M), adhesion (GO:7159 for genes EBI3, RAC2, BATF, CCL5, EOMES, CD3D, Cd7, LCP1, CD2, ITK, THEMIS, B2M), and migration (GO:2687 for genes SELL, RAC2, CCL4, CXCL13, CCL5, CXCL11, CXCL9, and CCL8). In contrast, group 110 primarily has terms associated with inflammatory response (GO:6954 for genes IL4R, MEFV, TGFB1, NLRP3, SIGLEC1, C5AR1, CD163, PIK3CG, NLRC4, TLR4, TNFRSF1B), apoptotic process (GO: 6915 for genes TGFB1, HGF, ABR, C5AR1, MKL1, NICAL1, MAP3K5, PIK3CG, INPP5D, NLRC4, TRAF1, TLR4, TNFRSF1B), and cell death (GO:43067 for genes STK10, HGF, ABR, SIGLEC1, C5AR1, MKL1, MICAL1, MAP3K5, PIK3CG, INPP5D, NLRC4, NCF2, TRAF1, TLR4, JAK3). Thus higher level clusters such as immune response are parsed at the zero layer according to specific biological sub-networks.

**Figure 5.**
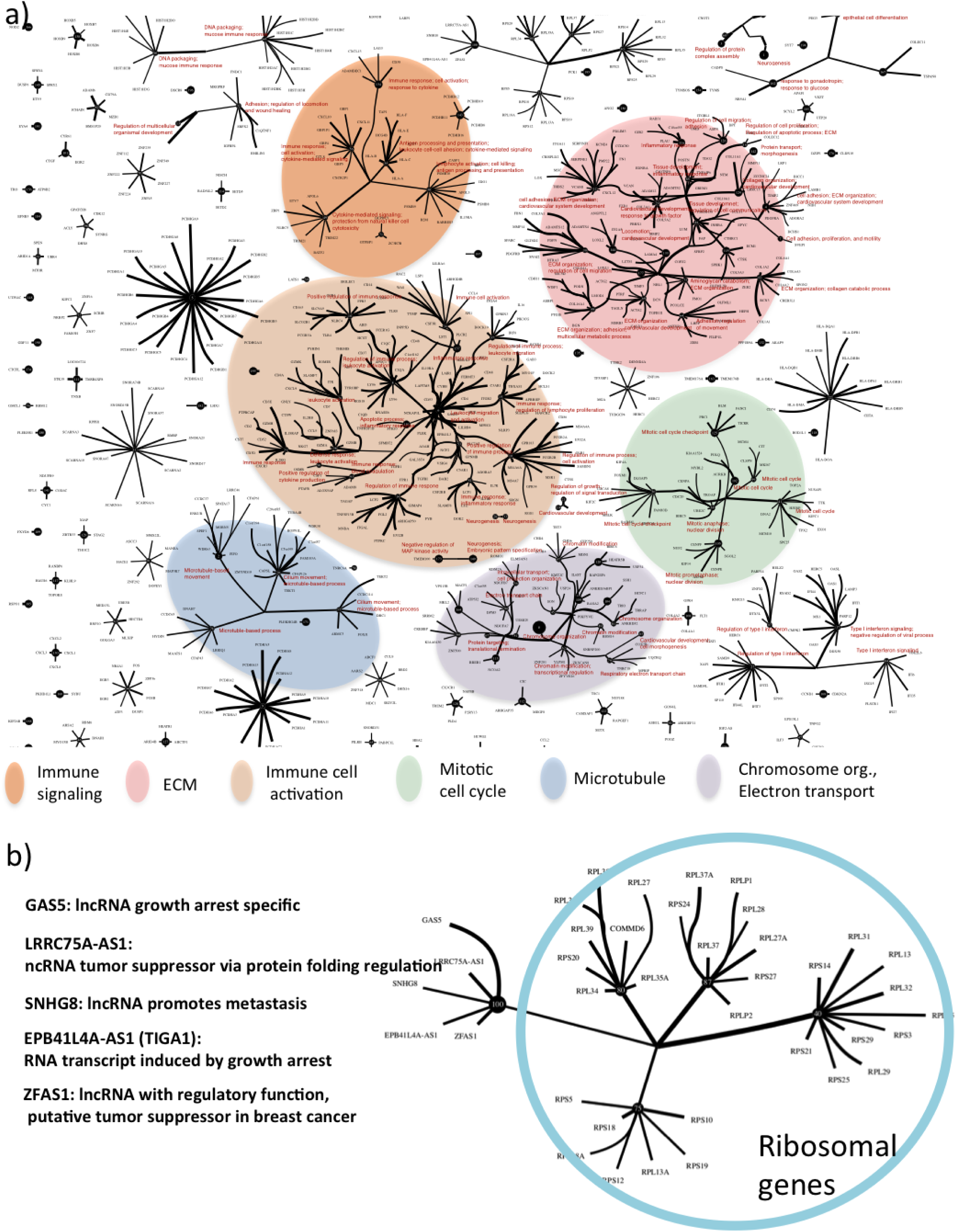
Gene function relationships in the latent factor hierarchy. a) Hierarchical factor relationships are shown exhibiting a tendency to cluster layer one groups with similar general Gene Ontology enrichments. Lines are shown connecting the top X strongest relationships and the top 1000 genes. Thicker lines indicate higher CorEx relationship probabilities. Representative GO terms for hierarchically clustered groups only are shown in red. Though this figure highlights clustering at layer one and above, many nonclustered layer one factors have very significant annotations as well, detailed in Fig. 2 and the supplementary information. b) In addition to general relationships, specific causal relationships are also detected at the upper layers. Here an example is shown in which noncoding RNAs that are implicated in tumor growth and metastasis via regulation of ribosome activity are isolated in a layer zero group and joined at the higher level to ribosomal structure genes. Noncoding RNA functional annotations are taken from the NCBI Gene database (http://www.ncbi.nlm.nih.gov/gene).

A particularly striking example of the kind of information that can be gleaned from the layer one factors is shown in Figure 5b. One can see a layer one factor (node) that joins four clusters of ribosomal proteins. The fifth layer zero cluster is comprised of a set of non-coding RNAs related to cancer metastasis through regulation of growth arrest and protein folding. Thus a cluster of non-coding regulatory factors is linked to the downstream interactions of proteins they influence. While the particular information highlighted here is already known, the presence of these meaningful relationships suggest that novel relationships can, in principle, be discovered by CorEx. This is more likely to happen when many thousands of samples are available to allow less common relationships and perhaps even tumor-specific rewiring of interactions to be seen.

### Patient stratification based upon combinations of latent factors yields differential survival associated with tumor-activated pathways

The continuous labels for the CorEx latent factors can be used as predictors of survival under a Cox proportional hazard model [26, 27]. A subset of the CorEx small to intermediate-sized gene groups was selected according to known biological significance (Gene Ontology enrichment) and used to stratify patients according to relative risk of death under a Cox proportional hazard model for the factor scores. We first constructed models for each factor individually. When this is done and patients are stratified roughly into thirds according to low, intermediate, and high relative risk of death, several gene expression groups appear to be associated with differential survival, as judged by the p-value for the difference between the two extremal Kaplan-Meier survival curves (Figure 6a and Supplementary Figure 6). False discovery rates (FDRs) were also estimated by randomly permuting the sample scores for each latent factor (shuffling each column) in the score matrix. The top single factor groups appear somewhat weak by this measure as the values are strongly limited by the small number of patient samples versus the number of potentially relevant networks. Using this analysis, the top latent factor group, related to immune chemokine and interferon signaling, yields an FDR level close to .3. Genes with large MI with respect to this group factor include among others CXCL10, CXCL11, CXCL9, CXCL13, ISG20, GBP (1-5), OASL, HERC5, STAT1, IFIT1, IFI35, TRIM22, and JAK2. Other groups that exhibit survival associations encompass additional immune system factors, fibroblast growth factor regulation, transforming growth factor beta signaling, map kinase regulation, Wnt signaling, Jak-STAT signaling, and cellular metabolism.

**Figure 6.**
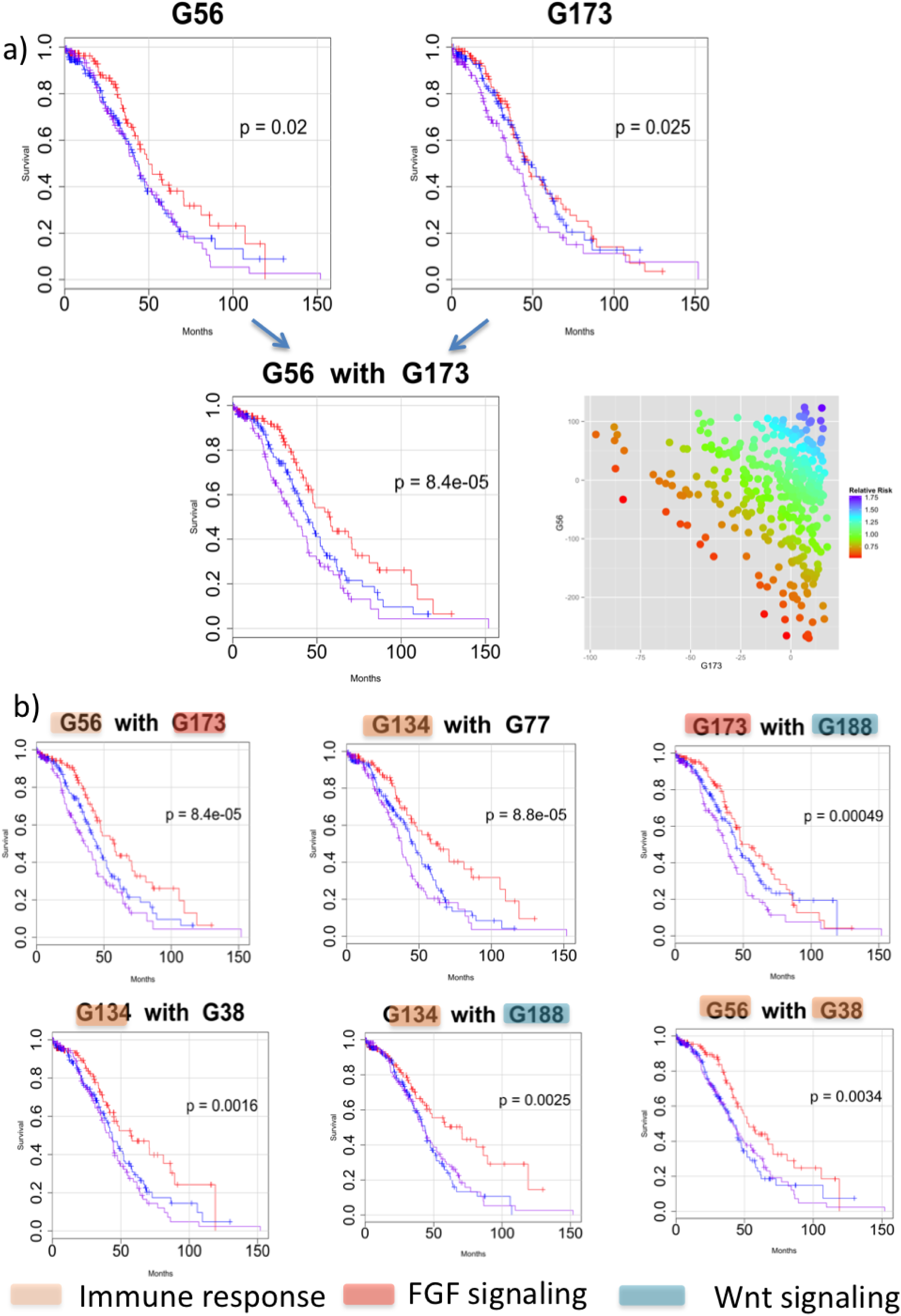
Group survival association and combinations. a) A Cox proportional hazard model is used to estimate relative risk for patients according to individual factor label values. Using this model, a relative risk of death can be calculated for each patient based upon her group label value. When patients are stratified into thirds according to relative risk, some factors show an association with survival indicated by the difference in Kaplan-Meier survival curves for the extremal groups (p-values shown in the graphs). The figure shows survival curves for two such factors, G56 and G173. The bottom 30%, middle 40%, and top 30% in terms of predicted survival under the coxph model are plotted in magenta, blue, and red, respectively. When these two factors are combined into a single risk model, the survival differential increases dramatically. b) Single factor combinations with various Gene Ontology annotations can be combined and show synergy relative to survival association. The false discovery rate under a randomization test for six such pairs is below about 1/3 compared to only one single factor achieving that level. Note that it is not suggested that factors be selected based solely upon survival associations but rather factors that are interesting for other reasons, e.g. they contain druggable targets, can be investigated for an impact on survival both alone and in combinations.

A perhaps even more compelling finding is that combinations of these statistically weak factors yield increased significance as pairs relative to a randomization test for paired factors, with 16/21 combinations yielding FDRs less than about .35 (Figure 6b). This suggests synergy of individual factors in terms of impact on survival. The pair of factors that show the greatest association with survival is immune system cytokine signaling combined with regulation of the fibroblast growth factor pathway. These two general features are hypothesized to be associated with long term survival in ovarian cancer from other studies, the latter due to its relationship to chemo resistance [28, 29]. The advantage here with respect to these previously known prognostic features is that each patient is assigned a relative risk according to activation of related pathways and therefore shows how they may combine to influence survival, indicating patients who might benefit from either single or simultaneous treatment with immune system modulation or FGF pathway inhibitors. Additionally, the fine parsing of overall immune system function by the sub-networks present in different groups has the potential to discriminate among different immunotherapies in terms of likely efficacy on a personalized basis. Overall, most of the top survival-associated combinations involve immune-related factors, emphasizing the primacy of the immunological milieu in influencing ovarian cancer survival rates in this cohort under the standard treatment protocols. The combination of a factor related to Wnt signaling and one for fibroblast growth factor signaling also appears predictive of survival in a subset of patients. Wnt signaling has also been associated with clinical prognosis in ovarian cancer [30]. Other factors and combinations of groups show narrower survival differentials. We expect that some of these play important roles for a relatively smaller proportion of patients and will gain significance once a greater number of tumor RNA-seq samples are available for analysis.

We performed survival analysis using PCA factors as well. While CorEx and PCA appear comparable in terms of the number of factors that can be used to stratify patients according to survival, the number and quality of biological enrichments for the CorEx factors is substantially greater than that for PCA (Figure 3b and Supplementary Figure 8).

It should be emphasized that in this retrospective analysis of a patient cohort that was treated prior to 2010, we are able to analyze the survival impact only with respect to standard carbo/taxol therapies as no targeted therapies were widely available or approved. Cell line and xenograft experiments or analysis of tissues from other clinical trials will likely bring to the fore other factor groupings that are associated with response to recently approved or experimental therapies, e.g. angiogenesis inhibitors, PARP inhibitors, and apoptosis signaling modulators to name a few [31, 32, 33].

### CorEx discovers differential expression in ovarian tumors of EMT-related genes previously implicated in breast cancer stemness and metastasis

A survival differential at longer times only is seen for one CorEx gene expression factor (denoted G159). This suggests that this factor may influence survival in a way that is not related to initial carbo/taxol response per se. Detailed examination of the Gene Ontology annotations for this latent factor cluster revealed genes associated with microRNAs in cancer (6 genes), chromatin modification (8 genes), regulation of transcription from RNA pol II promoter (10 genes), stem cell differentiation (5 genes), and regulation of Wnt signaling (5 genes). Further, when this factor is combined in a survival model along with the top single factor within the same CorEx run (G103, immune cytokine signaling), the resulting stratification yields a more significant survival association than any other pair of factors from that run (Figure 7a).

**Figure 7.**
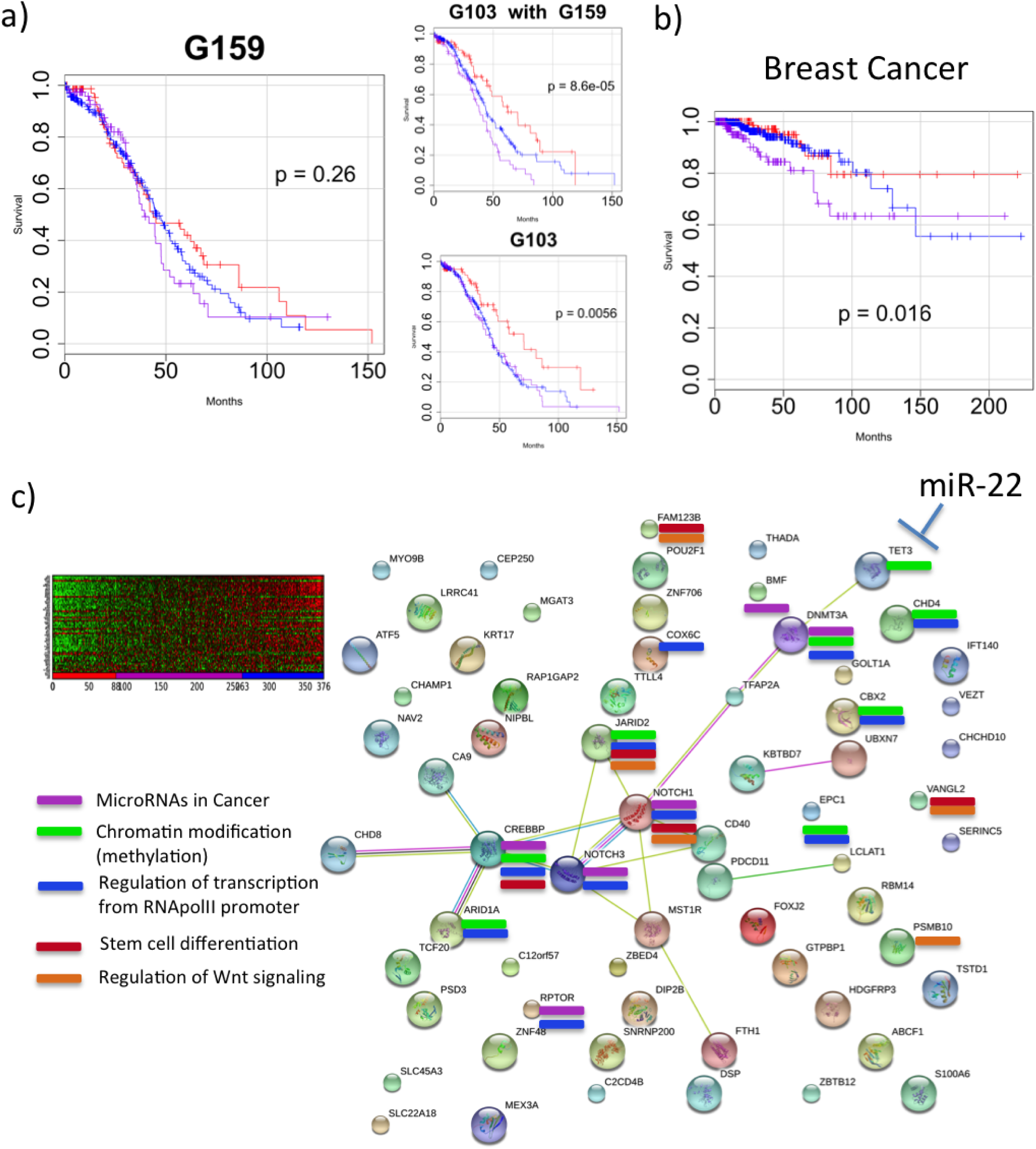
Discovery of new targets: regulatory microRNAs and EMT. a) One factor was observed to show a bias toward a difference in long-term survival. Though its survival differential was not significant alone, when put into a model combined with the factor for immune chemokine signaling, the resulting survival association was substantially greater and exceeded that of every other tested pair for that run.b) Breast cancer samples were mapped to CorEx factors learned from ovarian tumors and demonstrate a survival difference for this factor. c) The group contains a small network related to microRNAs in cancer, chromatin modification, and stem cell differentiation. Work in breast cancer associates the network with a cellular EMT phenotype and increased aggressiveness and metastasis of breast tumors. EMT features have been shown to be influenced through regulation by TET3 of the demethylase DNMT3A and represent a druggable target via the regulatory microRNA miR-22.

Intrigued by the possibility of a factor related to something more general than response to a specific chemotherapy, we sought to validate its significance using the TCGA breast cancer data. The reasoning behind this was that while breast tumors exhibit some similarity on a molecular basis to ovarian tumors, there are also significant differences. Earlier detection and a greater variety of chemotherapeutic options means survival will depend largely upon factors not directly related to carbo/taxol response. We calculated factor scores for TCGA breast cancer tumor RNA-seq samples across the factors determined by the ovarian cancer training samples (i.e. we did not train a new set of latent factors, but mapped the breast cancer samples to the factors learned from ovarian tumors). Using these scores to fit a stratified model with respect to the G159 latent factor yields a survival differential for breast cancer patients as well (Figure 7b). Subsequently we discovered that the genes in this group have important functions within a network that was recently studied by Song et al. [34] in preclinical models of breast cancer, where it was shown that TET3 controls expression of DNMT3A, a DNA methyl transferase that exerts specific influence on the chromatin state related to stemness (Figure 7c). Further, the network is implicated in the progression of breast cancer via the epithelial-mesenchymal transition, invasion, and metastasis. Mesenchymal-like properties of tumor cells and EMT-associated features in general have been associated with poor prognoses in ovarian cancer [35], though this particular network has not been previously implicated. Importantly, this network was shown to be controlled by a microRNA acting on TET3, thus providing a novel potential drug target [34]. Observation of this mechanism in ovarian cancer tumors suggests not only a new biological driver for recurrence but also the possibility for selection of ovarian cancer patients who might benefit from experimental miR-22 modulators being developed for breast cancer.

### Some latent factors can be used to distinguish between normal and cancerous ovarian tissue

We mapped RNA-seq profiles from normal ovarian tissue samples against the factors learned from the tumor samples and identified those that showed the greatest differences between the two types of samples. Many factors that exhibited large differences were found related to the mitotic cell cycle and replication, as would be expected when comparing cancerous to non-cancerous tissues. There were also several other factors that showed large distinctions and appeared related to a variety of other biological processes including ones for development and regulation of cell differentiation, neurogenesis, respiratory electron transport chain, sex differentiation, cell migration, and inflammatory response. Histograms of the factor scores for representatives of these groups are shown in Figure 8. These findings highlight some commonalities of pathway dysregulation among ovarian tumors. Further, several of the groups contain genes that can be targeted with existing drugs [36]. This presents the possibility that repurposing of existing drugs based upon this sort of data may provide new therapeutic options for a great many ovarian cancer patients, though the implications within these particular gene expression cohorts need to be further elucidated.

**Figure 8.**
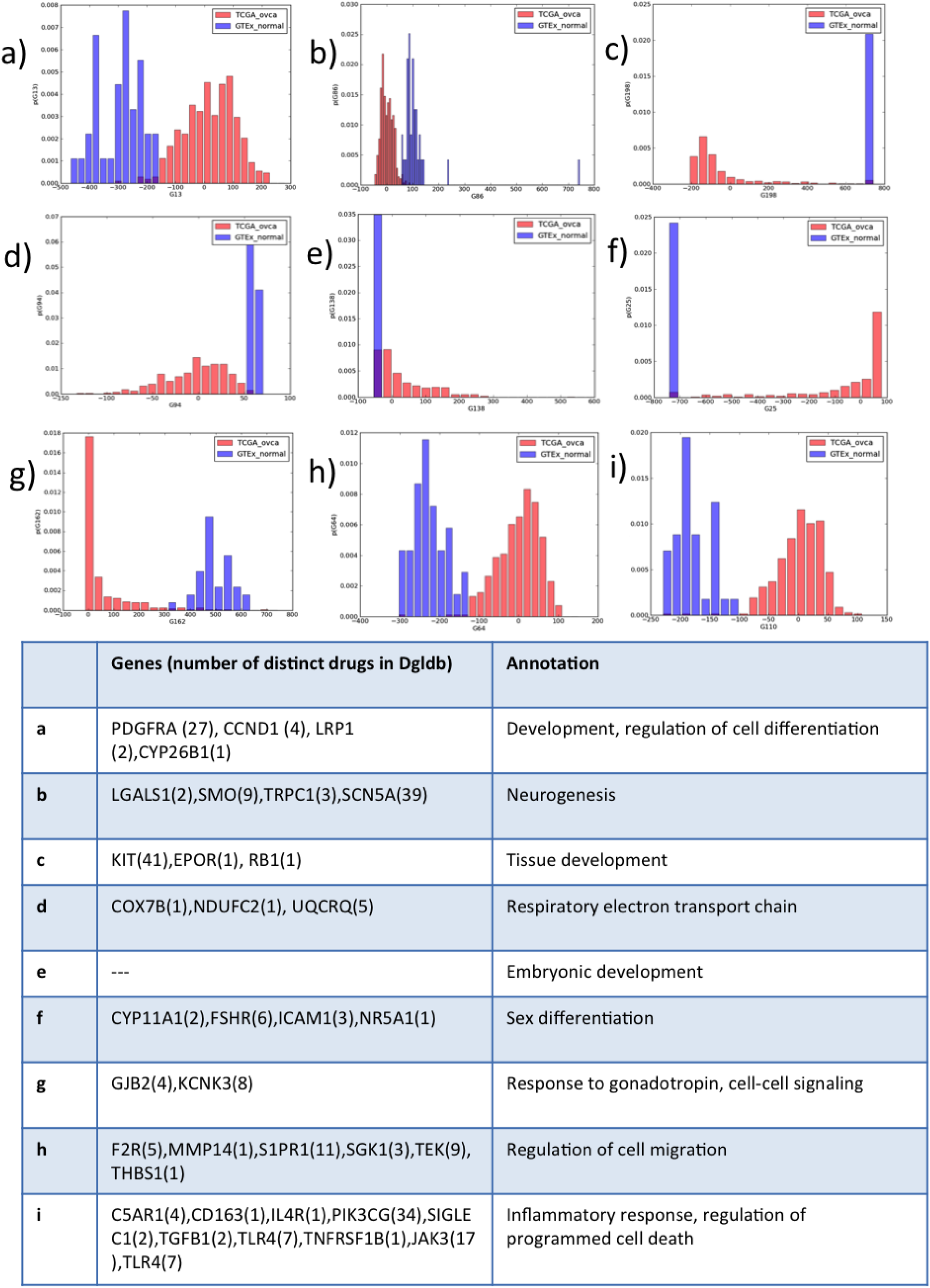
Comparison to normal ovarian tissue suggests druggable targets. When normal ovarian samples are assigned label values for the factors found from ovarian tumors, many exhibit striking differences. The histograms show factor scores in cases with large differences that correspond to novel drug targets in ovarian cancer (examples listed in Table 1).

## Discussion

In cancer and other diseases, high-throughput data combined with computational phenotyping i.e. the algorithmic distillation of observables that explain and allow prediction of the interaction of the course of disease and treatment, is poised to transform medicine as we know it. In this work we have presented a novel approach toward the goal of an RNA-seq-based computational phenotype. The method of correlation explanation leverages multivariate mutual information to infer complex hierarchical gene expression relationships directly from RNA-seq transcription levels, and the groups of genes thus obtained correspond to distinct cellular subnetworks as indicated by Gene Ontology annotations and known regulatory relationships. CorEx does this efficiently and without requiring any form of prior knowledge. The hierarchy of factors is shown to facilitate understanding and interpretability by relating networks at multiple levels. Though CorEx allows sharing of genes among groups, it also attempts to maximize overall independence between the groups in terms of expression patterns, with factors at higher levels being progressively less dependent. This leads to the ability to pick out groups from different upper level factors that may combine synergistically, and this is seen in the survival associations for some combinations of factors. In addition, tumor-specific assessment of differential transcriptional activation of the gene cohorts is intrinsic to the method. These results show how the observation of differential activity in these gene networks can have great clinical impact by enabling selection of patients most likely to benefit from particular therapies and/or combinations of therapies.

The use of RNA-seq, rather than microarray data, is essential for detailed inference of the transcriptional computational phenotype due to its superior dynamic range, sensitivity, and inherent whole transcriptome measurement capability compared to microarrays. Many past studies have focused on the use of microarray data, however RNA-seq should have substantial advantages for discovery of cancer-specific network interactions and this is supported by studies comparing the two [37, 38]. Some genes in the expression cohorts found here are not even measured in microarray experiments, often because their function is inadequately characterized and thus don’t justify inclusion of corresponding microarray probes. Also, uncommon transcript variants that arise from unstable cancer genomes cannot be detected with microarrays, resulting in misleading expression values. RNA-Seq data captures the effects of such alterations in ways that allow for their discovery and more accurate quantification [39, 40, 41]. The results presented here from such a relatively small patient cohort are extremely encouraging and clearly show that it is possible to extract a great deal of information from just a few hundred whole exome tumor transcriptomes and makes a strong case for obtaining larger numbers of samples in this context.

Early work on the clinical implications of large scale gene expression data in cancer included direct correlation of gene expression and responses of cell lines to cancer agents [42]. More recently, gene co-expression analyses based upon pairwise correlation or mutual information have been applied to infer networks and regulatory modules [24, 43, 44, 45], some of which are associated with cancer progression and prognosis. The number of modules found e.g. in breast cancer is markedly fewer (10-20 in the cited studies). Regulatory module inference in these cases often relies upon the application of graph theoretic techniques such as maximal clique identification to identify densely connected groups of genes, with special considerations to allow overlaps. CorEx, in contrast, constructs a graph with edges connecting multiple genes to latent factors rather than to one another and overlaps are intrinsic to the inference process. The approach in [46] is to define attractor metagenes using mutual information and, though relatively more restrictive in many ways and heuristic, is the most similar in spirit to CorEx. As far as we are aware, our work with CorEx demonstrates the first successful use of multivariate mutual information in the context of high throughput gene expression data. The principled approach of CorEx and also the addition of data-guided smoothing via the Bayes shrinkage prior allows for continuous improvement as more data is generated. These properties combined with favorable scaling suggest that CorEx can be widely adopted to analyze and interpret complex gene expression data in many contexts.

CorEx both provides a high level summary of the content of groups of differentially expressed genes and parses them out into particular groups for potential experimental and therapeutic purposes. Therefore it is an umbrella method that can encompass the particular findings of other studies on specific prognostic features of patient tumors. For instance, the original TCGA analysis of the ovarian tumor expression data used non-negative matrix factorization to identify a partitioning of genes and patients into a few subtypes [1]. Features of the four subtypes are clearly reflected in some of the higher level CorEx nodes such as that linking immune response factors. In contrast, here the functions broadly related to the putative subtypes have been parsed more finely on the lower level to reveal specific tumor-activated pathways and druggable targets. This allows one to make hypotheses regarding the interaction of functional gene modules in tumor development and progression. Further, the impact of expression modulation of such relatively smaller groups of genes may be more easily tested in cell lines and xenograft models.

There are many possible extensions and further applications of the methods described here. While mRNA provides a functional readout of many different causal processes, other types of salient information, e.g. somatic mutations, copy number variation, or chromatin marks related to gene activation or repression can be easily correlated with the gene expression groups at the higher level to directly link known biological causes to gene expression cohorts. This information can then point to specific mechanisms active in individual tumors or strengthen the RNA insights with the integration of complementary information. Though beyond the scope of this initial work, pan-cancer analysis is obviously of great interest, and CorEx’s favorable scaling behavior should allow for straightforward application in that context. Additionally, CorEx can be applied to other TCGA cancer types alone where one is likely to see different networks implicated.

## Conclusions

We have demonstrated a new technique to detect and comprehend the significance of coordinate gene expression in tumor subsamples. Due to the use of multivariate information, the algorithm is exceptionally sensitive to groups displaying multiple weak interactions. This property enables the discovery of an unprecedented number of functional biological expression groupings. In ovarian cancer, where there is a particular need for tools to aid in he development of both novel and rational combination therapies, we have shown that this method of analysis discovers new targets and lends itself to the selection of therapeutic combinations for individuals.

## Declarations

### Abbreviations

CorEx: Correlation Explanation
DNA: Deoxyribonucleic acid
EMT: Epithelial-mesenchymal transition
FDR: False discovery rate
GO: Gene Ontology
IC: Information content
GTEx: Genotype Tissue Expression
KEGG: Kyoto Encyclopedia of Genes and Genomes
MI: Mutual information
mRNA: Messenger ribonucleic acid
NCBI: National Center for Biotechnology Information
PARP: Poly ADP ribos polymerase
PCA: Principal component analysis
PPI: Protein-protein interaction
RAM: Random access memory
RNA: Ribonucleic acid
RPKM: Reads per kilobase of transcript per million mapped reads
TC: Total correlation
TCGA: The Cancer Genome Atlas

### Ethics and consent to participate

All human data analyzed in this study was originally obtained the TCGA Research Network and per their statement: “All specimens were obtained from patients with appropriate consent from the relevant institutional review board.”

### Consent to publish

No identifying details or images of individuals are included in this work.

### Competing interests

The authors declare that they have no competing interests.

### Author’s contributions

SP drafted the manuscript with assistance from GV. SP and GV conceived and designed the studies. GV developed the CorEx algorithm enhancements and ran it on the data. SP selected and prepared the training data and performed all analyses downstream of CorEx. All authors read and approved the final manuscript.

### Availability of data and materials

Supporting data is available in the supplementary files. The algorithm implementation used in this work can be obtained from GV or github (upon publication).

### Funding

GV acknowledges support from AFOSR grant FA9550-12-1-0417 and DARPA grant W911NF-12-1-0034 for development of the general algorithm used. The funders had no role in study design, data collection and analysis, decision to publish, or preparation of the manuscript.

## Additional Files

Additional file 1 – SupplementaryFiguresandText.pdf Supplementary methods details and text including a description of the Bayes prior and reproducibility across runs.

Additional file 2 – trainingmatrix.tcga ov.geneset1.log2.varnom.RPKM.xlsx Expression matrix used for training.

Additional file 3 – shrinkage2.groups.xlsx Group gene membership.

Additional file 4 – corex factor labels.xlsx Labels for each tumor sample.

## Footnotes

1 “Total correlation” is a historical misnomer, since it is not a measure of correlation, but of dependence.

